# *Cis*-regulatory elements of the cholinergic gene locus in the silkworm *Bombyx mori*

**DOI:** 10.1101/2021.03.29.437568

**Authors:** Kota Banzai, Susumu Izumi

**Affiliations:** Department of Biological Sciences, Kanagawa University, Hiratsuka, Kanagawa, Japan; Department of Cellular Biology, University of Georgia, Athens, GA, USA

**Keywords:** Cholinergic neuron, gene expression, *ChAT*, *Bombyx mori*

## Abstract

Genes of *choline acetyltransferase* (*ChAT*) and *vesicular acetylcholine transporter* (*VAChT*) are encoded in the same gene locus, called the cholinergic gene locus. They are essential in cholinergic neurons to maintain their functional phenotype. The genomic structure of the cholinergic gene locus is conserved among invertebrates to mammals. However, it is still inconclusive how cholinergic genes express only in cholinergic neurons in insects. In this study, we analyzed the upstream sequence of cholinergic gene locus in the silkworm *Bombyx mori* to identify specific *cis*-regulatory regions. We found multiple enhancer regions that are localized within 1 kb upstream of the cholinergic gene locus. The combination of promoter assays using small deletions and bioinformatic analysis among insect species illuminates two conserved sequences in the *cis*-regulatory region: TGACGTA and CCAAT, which are known as the cAMP response element and CAAT box, respectively. We found that dibutyryl-cAMP, an analog of cAMP, influences the expression of *ChAT*. Tissue-specific expression analysis of transcriptional factors showed potential candidates that control the cholinergic gene locus expression. Our investigation provides new insight into the regulation mechanism of cholinergic neuron-specific gene machinery in this lepidopteran insect.

## INTRODUCTION

Various types of neurons exist in the nervous system. The classification of each neuron is defined by its neurotransmitter phenotype; the expression of the specific genes encoding proteins for synthesis, storage, and release of neurotransmitters. Acetylcholine (ACh) is a major excitatory neurotransmitter in the central nervous system of insects (Gupta, 1987) and the neuromuscular junction in vertebrates (Krnjevic and Miledi, 1958). In insects, cholinergic neurons play a role in wing movements, locomotion, learning, and memory (Gauglitz and Pfluger, 2001, El Hassani et al., 2008). In cholinergic neurons, choline acetyltransferase (ChAT) and vesicular acetylcholine transporter (VAChT) have been well known as marker proteins because their molecular functions are essential for maintaining cholinergic phenotypes (Ichikawa et al., 1997). ChAT works as a catalytic enzyme of the ACh from precursor chemicals. VAChT functions to transport and store ACh into the synaptic vesicles (Song et al., 1997). Interestingly, their functions are completely different, yet they are expressed from the same cholinergic gene locus. In all animal species so far examined, *ChAT* and *VAChT* genes share exons that usually encode 5’-UTR. This suggests that the gene regulatory mechanism for the cholinergic gene locus is conserved among animal species. However, it is still not conclusive how cholinergic neurons express their own essential genes. To address this point, the elucidating the transcriptional factors that works for specifying neuronal identity is pivotal.

The gene regulation of the cholinergic gene locus has been well-studied in *Drosophila melanogaster* as a model system. Immunohistochemical assays and transgenic fly lines containing the cholinergic promoter fused to a GFP determined the distribution of cholinergic neurons in *Drosophila* nervous systems (Yasuyama and Salvaterra, 1999; Salvaterra and Kitamoto, 2001). The reporter gene analysis tested various upstream regions of the cholinergic gene locus and it indicated that the different sets of *cis*-regulatory elements worked for specific expression patterns in the nervous system (Kitamoto et al., 1992). The proximal region (−0.8 kb to -0.5 kb of the cholinergic upstream sequence) contained the necessary elements for *ChAT* expression in the central nervous system (CNS). The same region is sufficient to rescue the lethal phenotype of the *Drosophila Cha* mutant (Kitamoto and Salvaterra, 1993). POU-19/pdm-1 has been identified as a potential transcriptional factor to bind to a 22 bp sequence that is located in the proximal region (Kitamoto and Salvaterra, 1995). In contrast to the proximal regulatory region for the expression in CNS, the distal part of the upstream sequence is required for the expression in the peripheral nervous system (Kitamoto et al., 1992). Abnormal chemosensory jump 6 (Acj6), has been identified as a transcriptional factor that positively controls the *ChAT* mRNA expression in primary olfactory neurons (Lee and Salvaterra, 2002). Transcriptional regulatory mechanisms of the cholinergic gene locus have been previously studied. However, what transcriptional factor works for the CNS specific gene expression in insect species is still not completely understood.

Identification of distinct *cis*-regulatory elements is a major step in understanding the transcriptional mechanism regulating target genes. Classically, *in vivo* reporter assays using deletions or mutations in the promoter of a target gene is utilized to identify *cis*-regulatory elements. But such experiments have been inapplicable for non-model organisms like the silkworm due to the difficulty of making and maintaining transgenic lines. The baculovirus-introduction method was developed and adapted to study *cis*-regulatory elements in arthropods (Lu et al., 2005, Shiomi et al, 2003, Chen et al., 2020). This technology allows us to investigate the *in vivo* reporter assay in a non-model organism such as silkworm. Our previous report characterized the cholinergic gene locus in the silkworm, *Bombyx mori* (Banzai et al., 2015). We identified that the 2 kb upstream region of the cholinergic gene locus contained essential regulatory elements for expression in the CNS. In this paper, by the promoter assay based on the baculovirus-introduction method, we demonstrate that multiple *cis*-regulatory regions control the *ChAT* gene expression. Our investigation provides new insight into the regulatory mechanism underlying cholinergic neuron-specific gene machinery in the lepidopteran insect.

## RESULTS

### Promoter assay using sequential truncations illuminates the *cis*-regulatory regions

As reported in our previous paper (Banzai et al., 2015), the reporter protein, EGFP under the control of 2kb upstream region of *Bombyx* cholinergic gene locus expresses only in the CNS (Fig. 1B and 1B’), but not other tissues. Based on previous results, we constructed recombinant baculoviruses containing sequentially truncated promoters to investigate the cis-regulatory sequences (Fig. 1A). The EGFP signals were detected from the -2000 promoter to the -200 promoter (Fig. 1B-G), but not the -100, the 50, and the 0 promoters (Fig. 1H-J). The reporter signals from the sequentially truncated constructs decreased gradually depending on the length of the promoter region. Despite reducing the EGFP signals, the expression patterns seemed to be restricted to a specific subset of neurons. These results suggest that the upstream region between - 2000 promoter and -200 promoter functions to promoting expression level and does not regulate the expression specificity. We also quantified the activities of each promoter construct (Fig. 1K). The -2000 promoter to the -761 promoter showed a similar activity level. In contrast, the promoter activity decreased by approximately 40% in the -500 and the -300 promoters. In the -200 promoter, the promoter activities declined by about 65% compared to the -2000 promoter. The 100 and the -50 promoters exhibit similar levels to the 0 promoter, showing almost no promoter activity. These results from the sequentially truncated promoters indicate that three regions between -761 and -500, -300 and -200, - 200 and -100 are involved in the gene expression regulation of the *Bombyx* cholinergic gene locus.

**Figure 1.**
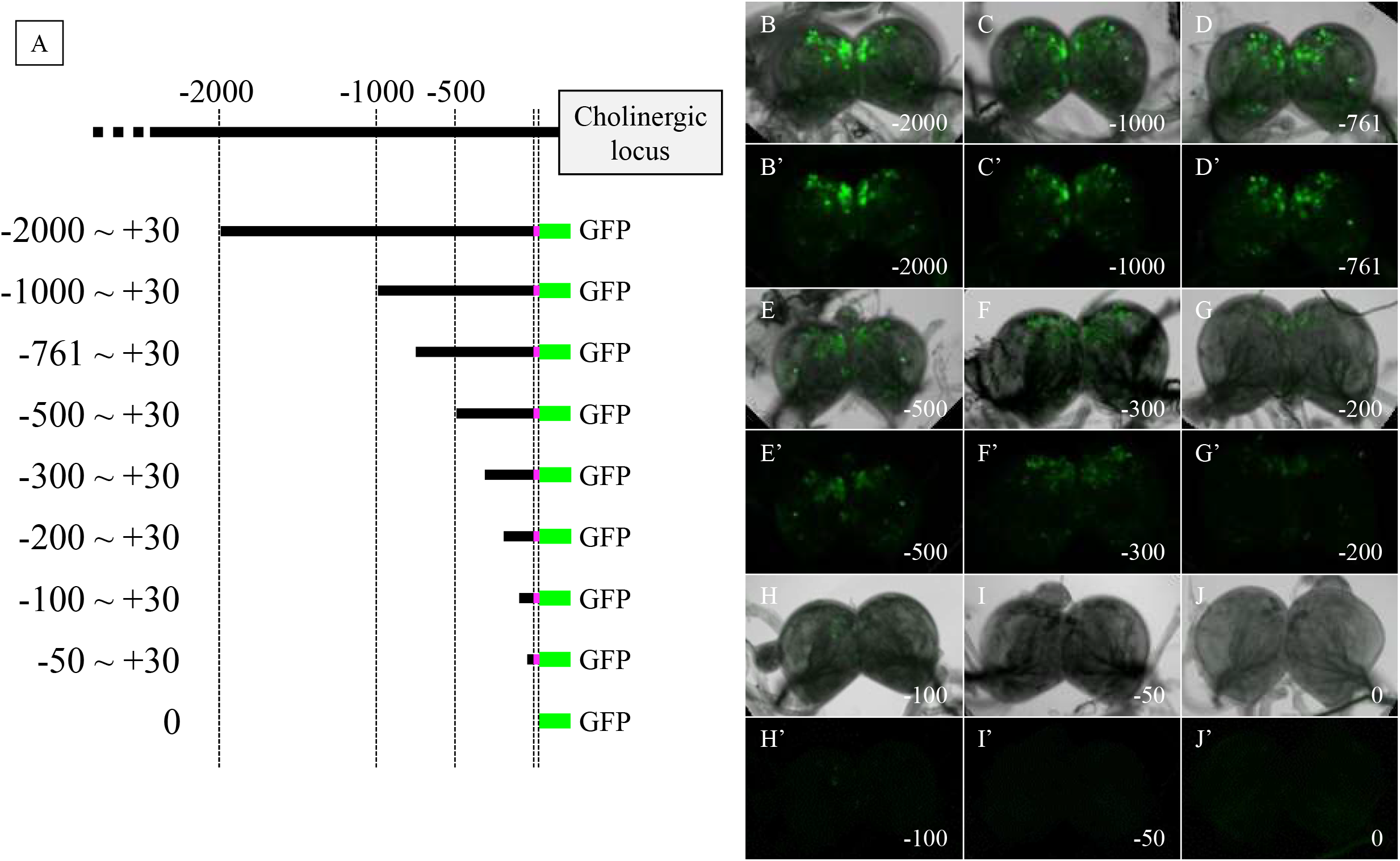
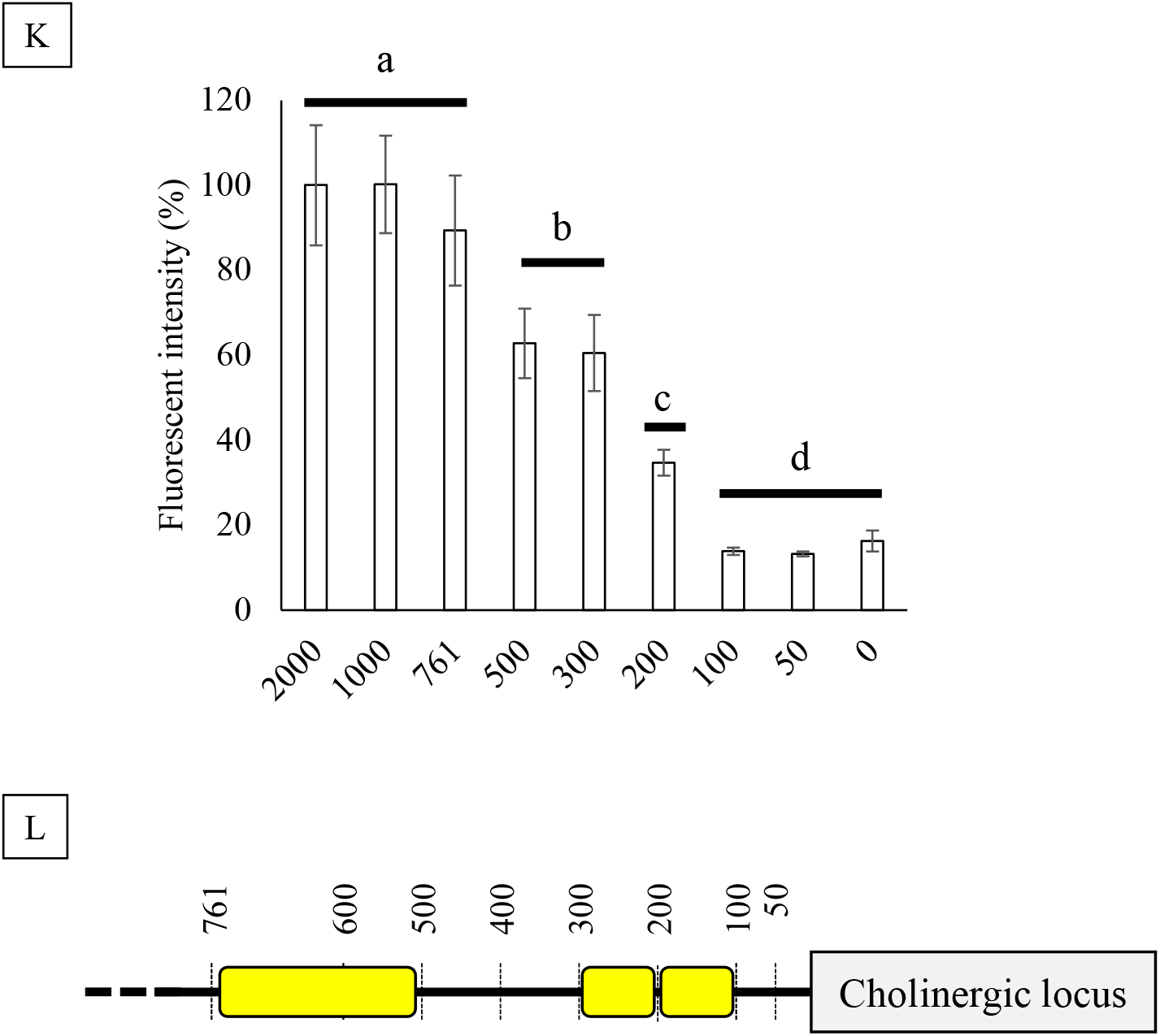
Promoter activity assays of sequential truncation promoters. (A) Schematic image of promoter constructs. Different length of ChAT promoter region is fused to EGFP protein. (B-J, B’-J’) EGFP expression patterns of different promoters in the larval brain. Pictures with alphabet were merged views with bright field and fluorescence. Pictures with alphabet plus (‘) were fluorescent views. Scale bars are 100 µm. (K) Quantitative fluorescent intensity. Data of each promoter activity are collected from multiple animals (n = 5). Small letters indicate significant differences (p < 0.05, ANOVA followed by the Tukey HSD test) among groups. If a non-significant difference was found between two groups, over-lapping letters are used to indicate the statistical significance. Each bar represents the mean ± standard deviation (SD) of assays. (L) The drawing diagram shows three regions (yellow boxes) identified via promoter assays.

### Conserved motifs among lepidopteran species

To find the DNA motifs that enhance the cholinergic gene expression, the identification of conserved sequences among different species is informative (ref). To do this among *B. mori* and other moths, we characterized the cholinergic gene loci from two lepidopteran species: the monarch butterfly, *Danaus plexippus*, and the postman butterfly, *Heliconius melpomene*, because their whole genomic sequence data have been published (Zhan et al. 2011, Heliconius Genome Consortium 2012). In the case of *D. plexippus*, putative *ChAT* and *VAChT* genes have been deposited in NCBI as OWR51509.1 and OWR51511.11, which are coding sequences. However, those two deposited genes don’t contain the shared regions, in which mRNAs of *ChAT* and *VAChT* have shared sequences in all species so far examined. This fact indicates that deposited sequences in *D. plexippus* didn’t cover the complete sequence of *ChAT* and *VAChT* inclusing 5’ and 3’ untranstared region (UTR). To determine the transcription initiation point of the cholinegic gene locus from *D. plexippus*, we identified the cholinergic gene exons of *D. plexippus* using the BLAST search using *Bombyx* exons as the query. We found the high similarity sequence (73.1%, 38/52) in the upstream region of *D. plexippus* OWR51509.1 against the first exon of *Bombyx* cholinergic gene locus (Fig. 2A and 2B). In the case of *H. melpomene*, we characterized its whole cholinergic gene locus by a similar procedure. The cholinergic gene locus of *H. melpomene* consists of 14 exons. As well as the *Bombyx* cholinergic gene locus, the first and second exons of *H. melpomene* encoded only 5’-UTR. The third exon encodes *VAChT*. Exons after the fourth exon are specific for the *ChAT* (Fig. 2A). The putative first exon has a high similarity (62.3%, 33/53) against the first exon of the *Bombyx* cholinergic gene locus (Fig. 2B).

**Figure 2.**
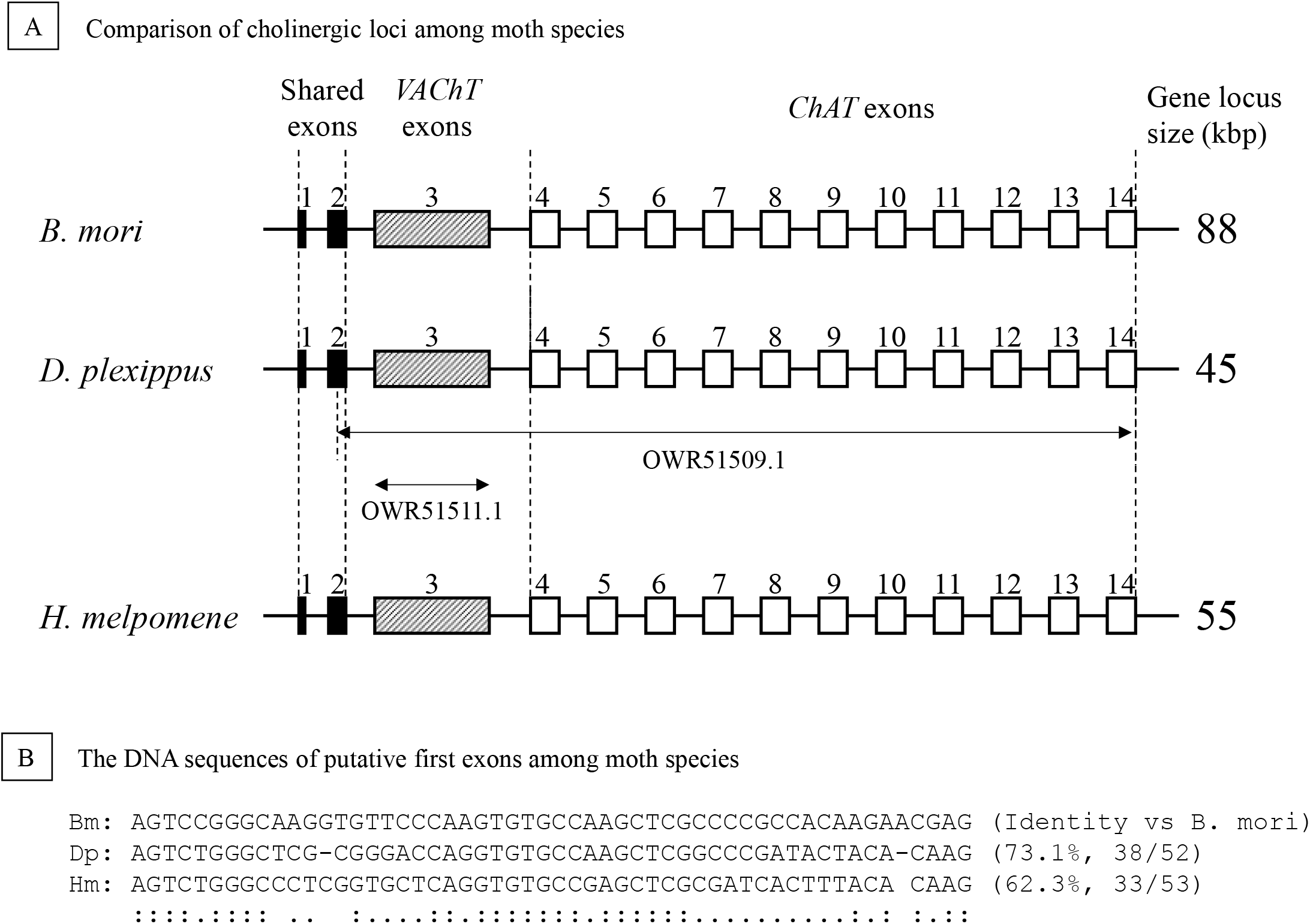
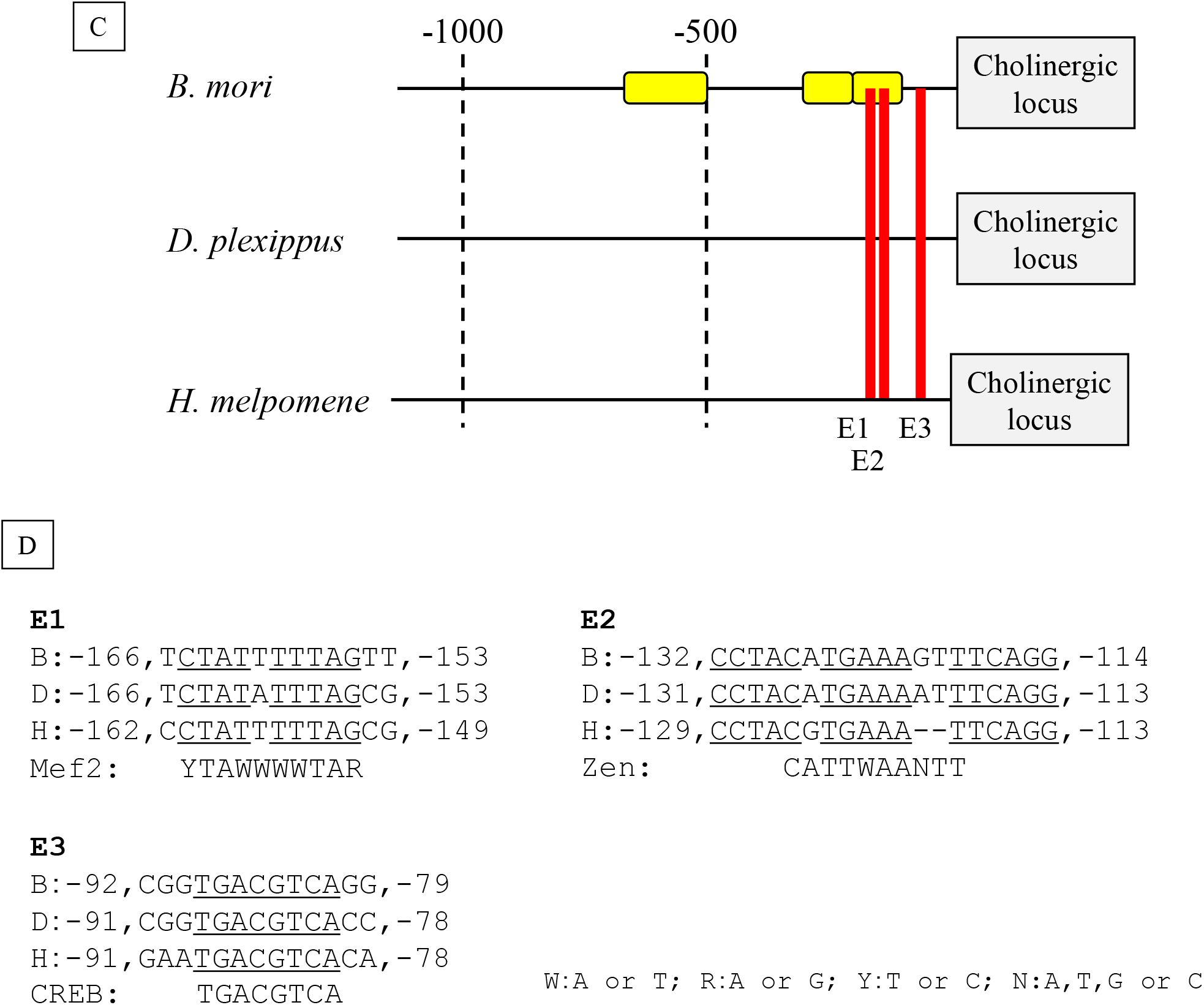
The cholinergic gene loci of three moth species. (A) Schematic representation of the cholinergic loci in *B. mori, D. plexippus* and *H. melpomene*. The genomic DNA is shown as a straight line. The exons common to *ChAT* and *VAChT*, specific for *VAChT*, and specific for *ChAT*, are indicated by black, striped, and white boxes, respectively. Approximate gene locus sizes (kbp) are shown on the right-hand side. (B) The DNA sequences of putative first exons among moth species. Abbreviations indicate each species; Bm: *B. mori*, Dp: *D. plexippus*, Hm: *H. melpomene*. Nucleotides conserved among three species and between two species are annotated by (:) and (.). The identity of nucleotide against *B. mori* sequence is shown on the right-hand side. (C) Schematic diagram of mussagl analysis. Three yellow boxes indicate the identified region via promoter assay in Fig. 1. Red bars represent calculated sequences with high similarity among moth species (E1, E2, and E3). (D) Nucleotide fragments are identified in (C). Each fragment possesses a potential consensus sequence, Mef2 (YTAWWWWTAR), zen (CATTWAANTT), and CREB (TGACGTCA), respectively.

Next, we compared upstream DNA sequences of these three lepidopteran species to find conserved sequences that had the potential to regulate gene expression. For this comparison, we used the Mussa program that visualizes conserved regions across multiple species. Our promoter assay indicated that the 1 kb upstream region of the *Bombyx* cholinergic gene locus is sufficient to control gene expression specifically in the CNS. We therefore selected the 1 kb upstream sequences of three lepidopteran species for this comparison. We found three conserved three motifs located within -200 bp upstream of cholinergic gene loci (E1, E2, and E3. Fig. 2C). To predict possible transcriptional factors that bind to the conserved sequence, we investigated three conserved sequences by the STAMP programs. As shown in Fig. 2D, the most distal element E1 contains a consensus sequence (YTAWWWWTAR) of Myocyte-specific enhancer factor 2A (Mef2A). The center element E2 contains a consensus sequence (CATTWAANTT) of zerknüllt (zen). The most proximal E3 contains a consensus sequence (TGACGTCA) of cAMP response element-binding protein (CREB).

### Promoter assay using small deletions reveals *cis*-regulatory elements

To validate whether the conserved sequences are required for the gene expression of *Bombyx* cholinergic gene locus, we tested promoter activities by introducing small deletions into the promoter sequence. As shown in Fig. 3A, the deletion between -200 to -150, including the E1 element causes a 65% reduction of the GFP signal compared to the -2000 promoter. Moreover, the deletion between -200 to -100, including E1 and E2 elements, causes a more reduction than a single deletion of the E1 element. These results suggest that E1 and E2 elements work together to regulate gene expression.

**Figure 3.**
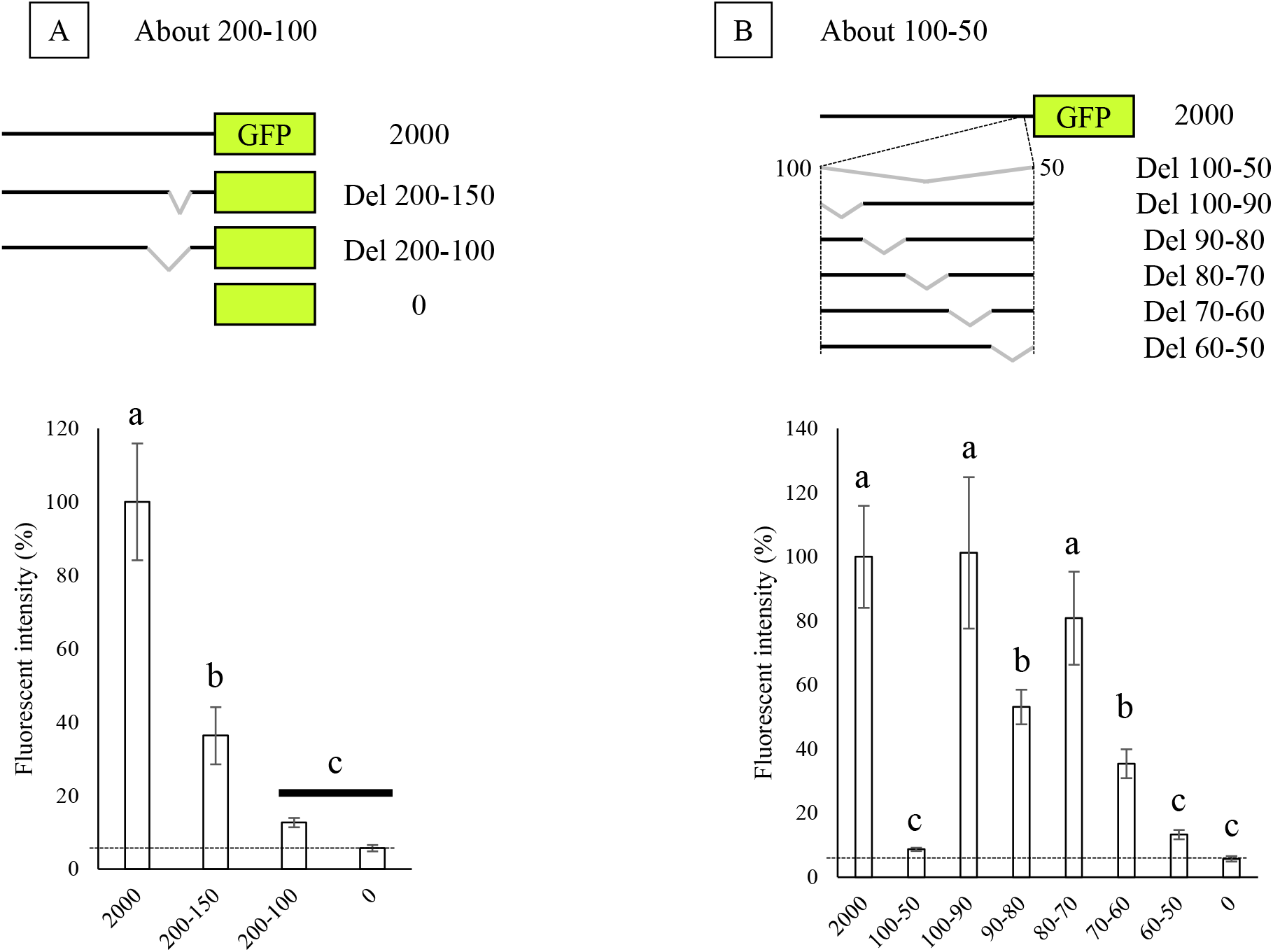
Promoter activity assays of deletion promotes. Quantitative fluorescent intensity using deletions spanning -200 to -100 (A) and -100 to -50 (B). Data of each promoter activity are collected from multiple animals (n = 5). Small letters indicate significant differences (p < 0.05, ANOVA followed by the Tukey HSD test) among groups. Each bar represents the mean ± standard deviation (SD) of assays.

Next, we analyzed the promoter activity of the E3 element which is located most proximally. A deletion promoter, del 100-50, showed a 90% reduction of promoter activity (Fig. 3B). To narrow down the critical sequence for the gene expression regulation, we deleted DNA sequences in every 10 bp between 100 bp and 50 bp of the promoter region. As shown in Fig. 3B, the del 90-80 promoter excluding the E3 element shows a 45% reduction of promoter activity. This result supports the idea that the consensus sequence (TGACGTCA) of CREB has an important role in the *ChAT* gene expression. Additionally, two deletion promoters, del 70-60 and del 60-50 caused drastic reductions of promoter activity, 65% and 90%, respectively (Fig. 3B). However, between -60 to -50, we could not observe any preserved motifs among lepidopteran species.

### cAMP modulates the expresion level of *Bombyx ChAT* gene

Our promoter assay using small deletion mutants revealed that the promoter region -100 to -50 consists of three different elements. We predict that the identified regions should be conserved not only moth but also other insect species if they are critical for gene expression. Thus, we expanded the bioinformatic comparison of the promoter region among multiple insect species. As shown in Fig. 4A, the CRE sequence (TGACGTCA) in -90 to -80 and a five-nucleotide sequence (CCAAT) in -70 to -60 were conserved among seven insect species. Interestingly, the distance between the CRE sequence and the CCAAT was withing 21 nucleotides of each other in these six insect species (Fig. 4A). This indicates us that the combination of two elements work synergistically. It has been reported that CREB forms heterodimers with C/EBP (CCAAT-enhancer-binding protein) *in vitro* (Vinson et al., 1993). We could not find any conserved motif in the most proximal part (−60 to -50).

**Figure 4.**
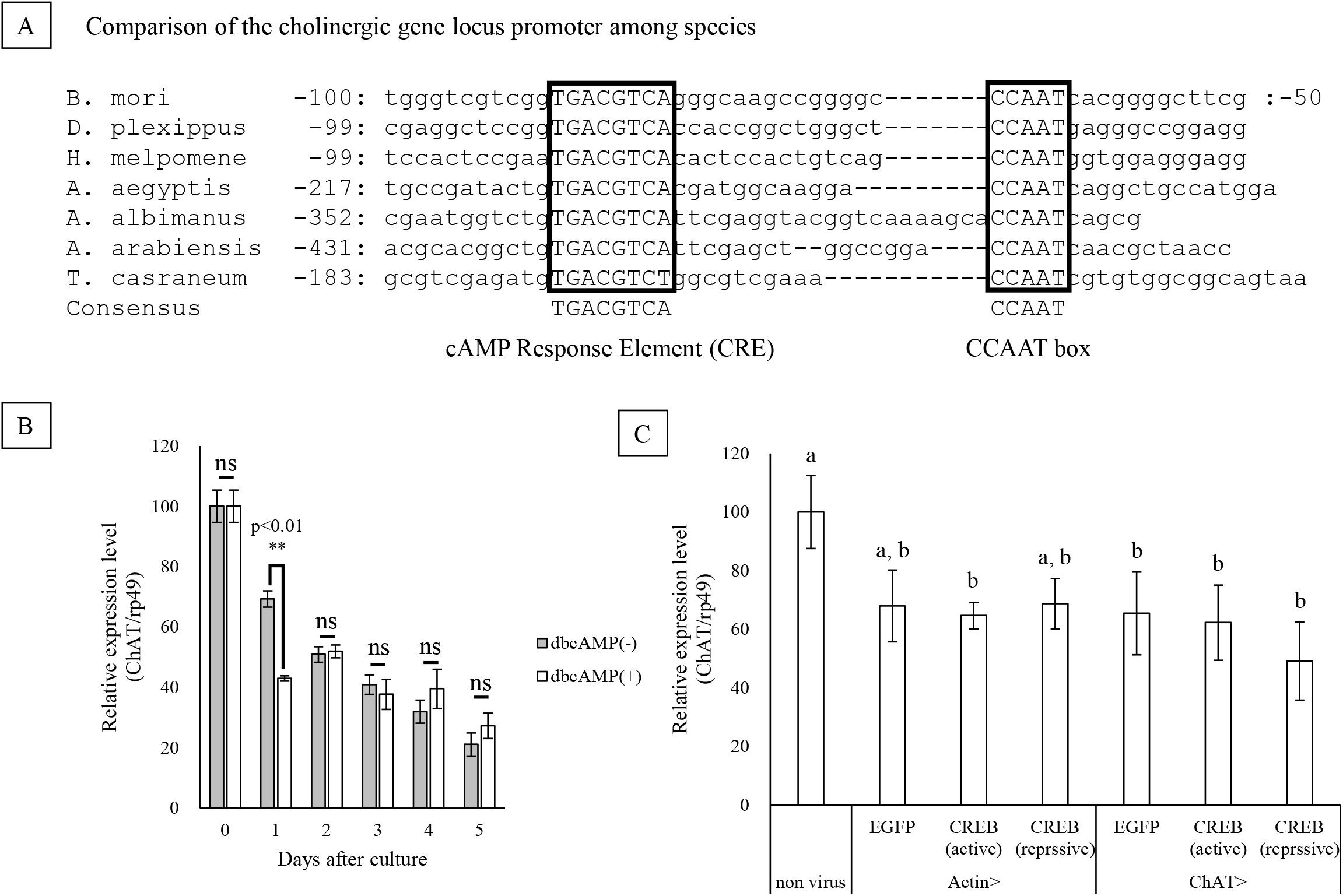
dbcAMP influences to the ChAT gene expression. (A) Comparison of the cholinergic gene locus promoter among insect species. Nucleotide sequences are collected through the Blast using the genome database of each insect species. Two conserved sequences, TGACGTCA and CCAAT are surrounded by boxes. The *ChAT* expression levels in (B) and (C) were stabilized by a housekeeping gene, *rp49*. Data of each condition are collected from multiple animals (n = 3). Each bar represents the mean ± standard deviation (SD) of assays. (B) qPCR analysis in *ex-vivo* cultured larval brain between dbcAMP (−) and (+). Asterisk (*) indicates significant differences (p < 0.01, ANOVA followed by the Tukey HSD test). (C) qPCR analysis in larval brain injected actCREB or repCREB virus vectors. Small letters indicate significant differences (p < 0.05, ANOVA followed by the Tukey HSD test). If a non-significant difference was found between two groups, over-lapping letters are used to indicate the statistical significance.

We decided to focus on CREB based on our promoter assay and the bioinformatic analysis. Cytoplasmic cAMP works as a second messenger, which activates CREB via the protein kinase A (PKA) signaling pathway (Alberini, 2009). A previous study demonstrated that the addition of analog of cAMP, dibutyryl cAMP (dbcAMP), in mammalian PC12 cells (naturally expresses both ChAT and VAChT) increases the amount of ChAT activity (Shimojo et al., 1998). Thus, we examined the effect of dbcAMP on the *Bombyx ChAT* gene expression in the *ex vivo* brain. As shown in Fig. 4B, the different expression level of the *ChAT* gene between brains cultured with or without dbcAMP was observed 24 hours after starting culture, but not other time points. This result indicates that cAMP works negatively for *Bombyx ChAT* gene expression. However, this result is not consistent with the promoter assay that we predicted that CREB, a downstream transcriptional factor of cAMP response, works as a positive enhancer (Fig. 3B). To test the function of CREB in *ChAT* gene expression more directly, we designed the experiment that we express active and repressive forms of CREB in larvae through recombinant baculoviruses and measure the *ChAT* gene expression by quantitative PCR. It is well known that serin 133 (S133) in human CREB gets phosphorylated by cAMP stimulation and the phosphorylation is required for the interaction with co-transcriptional factors (Gonzalez and Montminy, 1989, Suzuki et al., 2011). The mutation into the amino acid sequences including S133, RRPSYR to DIEDML, makes CREB proteins constitutively active (Cardinaux et al., 2000). In *Drosophila*, an artificially modified CREB that lacks the leucine zipper domain functions as dominant-negative (Yin et al., 1994). We expressed active and repressive forms of CREB ubiquitously by Actin promoter or ChAT-positive neurons by the ChAT promoter. As shown in Fig. 4C, *ChAT* gene expressions in control, active or repressive forms of CREB were comparable level. This data suggests that CREB is not a regulatory protein for this locus, and other transcriptional factors are involved in the expression of cholinergic gene locus.

### Gene expression profile of candidate transcriptional factors

We narrowed down the candidates for activation of *ChAT* expression using the STAMP program to uncover which transcriptional factors may bind the upstream region of the *Bombyx* cholinergic gene locus. We selected three regions (−761 to -500, -300 to -200, and -200 to -100) identified in our promoter activity assays. As summarized in Fig. 5A, the STMAP program showed sixteen genes for -761_-500, six genes for -300_-200, and two genes for -200 _-100. We tested the expression pattern of those predicted transcriptional factors in six tissues. Fifteen genes out of the candidate twenty genes were expressed in the brain (Fig. 5B). Moreover, four genes, *deformed* (*Dfd*), *even-skipped* (*eve*) *tailless* (*tll*), and *twin of eyeless* (*toy*), were expressed in the brain and not in any other tissues.

**Figure 5.**
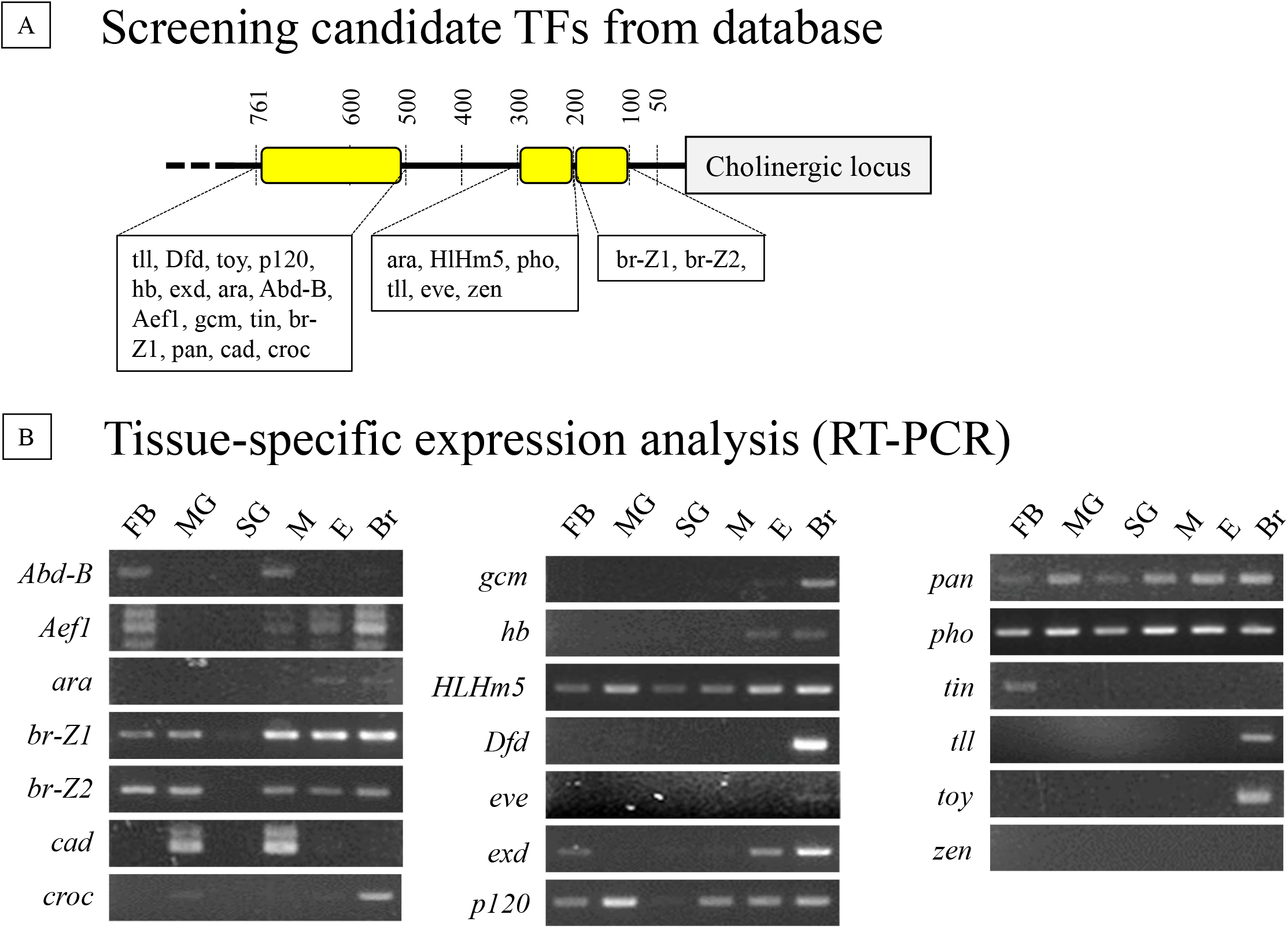
Gene expression patterns of candidate transcriptional factors. (A) The schematic structure of the cholinergic gene promoter region. Yellow boxes indicate that three identified regions in Fig. 1. Candidate transcriptional factors predicted by STAMP are described. Detail data of E-values against each TF consensus sequence is listed in Supplementary Table 3. (B) Gene expression profile of each TF gene in different tissues; FB: Fat body, MG: Malpighian tube, SG: Silk grand, M: Midgut, E: Epidermal cells, Br: Brain. Primer information of target genes is listed in Supplementary Table 4.

## DISCUSSION

We discovered four *cis*-regulatory elements on the *ChAT* gene promoter region. The most distal region spans from -761 to -500. We could not find any significant binding site based on the bioinformatic analysis. Further promoter assays using different sets of truncations and mutations of promoter sequences is required to discover regulatory elements located in this region. We found consensus sequences for the other three regulatory elements (named as E1, E2, and E3, Fig. 2C). The E1 element has a Mef2 consensus sequence. Mef2 expresses strongly in the mushroom body from embryo to adult in the fruit fly (Crittenden et al., 2018). Besides, *Mef2* is annotated as one of the enriched genes in insulin-producing cells (IPCs) in *Drosophila* (Cao et al., 2014). Previously, we identified that *ChAT* was expressed in the majority of the CNS including mushroom body and IPCs (Banzai et al., 2015). The overlapped expression pattern of *Mef2* and *ChAT* in CNS supports that Mef2 has a strong potential to regulate cholinergic gene expression. The E2 element has zen motif, but our gene expression analysis of several tissue samples didn’t show the expression of *zen* in the brain. The E3 element possesses a CRE motif where CREB transcriptional factor binds to. We tested the effect of CREB on *ChAT* gene expression by overexpression of activated or inactivated proteins. However, we did not obtain evidence that CREB influences *ChAT* gene expression. There are several methods to isolate sequence-specific binding proteins (e.g., the yeast one-hybrid assay, the electrophoretic mobility shift assays (EMSAs), and the reverse ChIP). In the future, we will apply those approaches to elucidate the regulatory proteins.

Previous reports in *Drosophila* have uncovered that -0.8 kb to -0.5 kb upstream sequence of ChAT start codon is an important regulatory region for *ChAT* gene expression in CNS (Kitamoto and Salvaterra, 1995). In addition, their DNase-I foot-printing analysis of the identified 0.3 kb revealed a 22 bp sequence that is conserved among *Drosophila* species and contains an octamer-like motif (ATTCAAAT). We analyzed but could not find the *Drosophila* identified sequence in *B. mori* and other insect species. Conversely, in *Drosophila*, we could not see the combination element of the Cre (TGACGTCA) and the CCAAT sequences observed in our study. Further advanced and optimized searching tools for transcriptional factor binding sites would help to resolve this issue.

*ChAT* and *VAChT* genes are essential in cholinergic neurons. However, there are other specific genes to maintain cholinergic neuronal identity. For example, acetylcholine esterase (AChE) breaks down acetylcholine to choline and acetic acid at the synaptic cleft to tightly control the signal transduction. The choline transporter (ChT) works to re-uptake the decomposed choline to presynaptic neurons, which is critical to resynthesize acetylcholine. Acetylcholine receptors (AChRs) are responsible for the autoregulation of neurotransmitter release. A previous report using *C. elegans* has identified a transcriptional factor, Unc-3 as a terminal selector for cholinergic motor neurons by analyzing promoters from several cholinergic neuron identity genes (Kratsios et al., 2012). Similar approach in *B. mori* would be crucial to elucidate the specific regulatory machinery in the future.

### Experimental procedures

#### Insects and cell lines

Eggs of the multivoltine strain, N4, of *B. mori* were obtained from Kyushu University (Fukuoka, Japan). Larvae were reared on an artificial diet, Silkmate II (Nihon Nosan Kogyo, Yokohama, Japan) at 25 °C under a 14-h light/10-h dark photoperiod cycle. The cell line of *Spodoptera frugiperda* Sf9 was maintained in SF900-III medium (Invitrogen) at 27 °C in a cell culture incubator.

#### Construction of the reporter plasmids

Different DNA fragments of the cholinergic gene locus promoters and *EGFP* ORF sequence were amplified using specific primers (Supplementary Table 1) and genomic DNA or pEGFP-1 (Clontech) as template DNA. The PCR products were inserted into a pFastBac-HTa vector (Invitrogen). The deletion plasmids were amplified by inverse PCR using specific primers (Supplementary Table 1) and circularized by self-ligation. We used KOD-FX neo (Toyobo) as DNA polymerase in our PCR reactions.

#### Preparation of recombinant AcNPVs and infection into larvae

Recombinant AcNPVs were prepared according to the instruction manual of the Bac-to-Bac baculovirus expression system (Invitrogen). The recombinant baculoviruses were amplified by infecting sf9 cells 2-3 times. The virus titer was determined by plaque assay (Shirata et al., 1999). The virus solution was diluted to 1-2 × 10^6^ pfu/ml with sf900-III medium before use. Virus injection and observation of samples are followed from a previous report (Banzai et al., 2015). We used multiple animals (n = 5) for measuring the promoter activity.

#### Determination of putative cholinergic gene loci from insect species

We utilized two moths (*Danaus plexippus* and *Heliconius melpomene*), three mosquitoes (*Aedes aegypti, Anopheles albimanus* and *Anopheles arabiensis*), and one beetle (*Tribolium castaneum*). Genome DNA sequences of insect species were obtained from online databases (Supplementary Table 2). We searched the cholinergic gene loci by BLAST of each database using data of *B. mori* cholinergic locus as the query.

#### Computer-assisted retrieval of the conserved DNA regions

The conserved nucleotide sequences among three lepidopteran species were predicted by comparing their upstream regions of the cholinergic loci using the Mussa program (http://mussa.caltech.edu/mussa) with changing parameters of the threshold (nt) and window (nt). The consensus sequences were searched from the conserved sequences by using the STAMP (Mahony and Benos, 2007, http://www.benoslab.pitt.edu/stamp/).

#### Isolation of total RNA, Reverse transcription, and PCR

We prepared total RNAs of 6 tissues: fat body, midgut, salivary gland, malpighian tubule, epidermal cells, and brain from newly ecdysed and non-fed fifth instar larvae. We collected each tissue from at least three individual animals. Total RNA was isolated following the acid guanidinium thiocyanate-phenol-chloroform extraction method (Chomczynski and Sacchi, 1987). The total RNA (200 ng) was converted to cDNAs by the PrimeScript™ RT reagent kit with gDNA Eraser (Takara Bio) in 10-µl reaction volume. The synthesized cDNAs were diluted with 90 µl of MilliQ water (final concentration of total RNAs is 2 ng/µl).

For the PCRs, the reactions were carried out in 20-µl reaction volumes containing 0.5 µl of template cDNAs (equivalent to 1 ng of total RNA), 1×GoTaq Green Master Mix (Promega), and primers at final 0.5 µM concentration. The PCR products were separated using a 2.0% agarose gel electrophoresis and stained with ethidium bromide. Information about the primers, annealing temperatures and the number of cycles for PCR reactions is summerized in Supplementary Table 4. The primer sequence for *pho* gene was obtained from Li et al. (2012).

#### Effect of cAMP on the *ChAT* gene expression

We dissected out the brains from newly-ecdysed and non-fed fifth instar larvae. Those brains were washed four times with PBS (20 mM NaPB, 150 mM NaCl, pH 7.0) and two times with TC100-insect medium (Sigma Aldrich). Then, the dissected brains were stored for one hour in a TC100-insect medium containing 10% fetal bovine serum and penicillinG/streptomycin cocktail (Roche). We added the dbcAMP (Sigma Aldrich) with a final 1% consentration into the medium. The tissues were incubated at 27 °C in a cell culture incubator. We collected five cultured brains at 24, 48, 72, 96, and 120 hours after the addition of dbcAMP. For each time point, we prepared three replicates. We operated all procedures in the culture hood to prevent contamination of bacterium except the brain dissection from larvae. Total RNA isolation and cDNA synthesis were performed by the same protocol as described above.

For quantitative PCRs (qPCRs), we used THUNDERBIRD™ SYBR® qPCR Mix (Toyobo), following the manufacturer’s instruction. The 20-µl reaction system contained six components: (1) 5 µl of cDNA (equivalent to 2 ng of total RNA), (2) 10 µl of THUNDERBIRD SYBR qPCR Mix, (3) 0.4 µl of 50× ROX reference dye, (4) 3.4 µl of SDW, (5) 0.6 µl of forward primer (10 µM), and (6) 0.6 µl of reverse primer (10 µM). The real-time PCR program was performed using 7300 Real-Time PCR System (Applied Biosystems) following the instrument manual. It consisted of an initial step at 95°C for 60 s, followed by 40 cycles of 95°C for 10 s and 60°C for 50 s. qRT-PCR quantitative assays of all samples were repeated three times. Primers used in PCR reactions were listed in Supplementary Table 1.

Effect of active or repressive CREB overexpression on the *ChAT* gene expression Total RNAs were extracted from larval brains by using Isogen (Nippon). cDNA was synthesized from the total RNAs by using the PrimeScrip RT reagent Kit with gDNA Eraser (Takara bio). DNA fragments of the CREB ORF and the repressive CREB (repCREB) were amplified using specific primers (Supplementary Table 1) and the brain cDNA and cloned into pFastBac-HTa vector. The conducted plasmids were named as CREB vector and repCREB vector, respectively. Next, we constructed a plasmid for the active CREB (actCREB) by performing the inverse PCR using specific primers (Supplementary Table 1) and the CREB vector. This vector is named as actCREB vector. The DNA fragment of the Actin A3 gene promoter (−127_+542 bp, GeneBank: U49854.1) and cholinergic gene locus promoter (−2000_+30 bp) are amplified by using specific primers (Supplementary Table 1) and genomic DNA. The PCR products were inserted into both of actCREB or repCREB vectors.

We prepared and injected the recombinant virus into animals with the same protocol described above. Five days after injection, we dissected out brains from the virus-infected larvae. We collected three brains from each experimental condition. We prepared three replicates for measuring *ChAT* gene expression by qPCR. Total RNA isolation, cDNA synthesis and the qPCRs were performed by same protocol as described above.

#### Selection of candidate transcriptional factors

We identified candidates of transcriptional factors that may be involved in expression regulation of cholinergic gene locus by using the STAMP (Mahony and Benos, 2007, http://www.benoslab.pitt.edu/stamp/). We individually characterized the four enhancer elements (−761_-500, -300_-200, -200_-100 and -100_-50) by using two datasets: flyreg (Bergman/Polland) and Fly (curated by Bergman).

### Statistical analysis

All data in graphs were shown as mean ± standard deviation (SD). Significant differences (p < 0.05) among multiple sample groups or conditions were analyzed using one-way ANOVA followed by the Tukey HSD test. Same letter indicates no significant difference. Different letters indicate significant differences. If a non-significant difference was found between two groups, over-lapping letters are used to indicate the statistical significance.

**Table 1.**
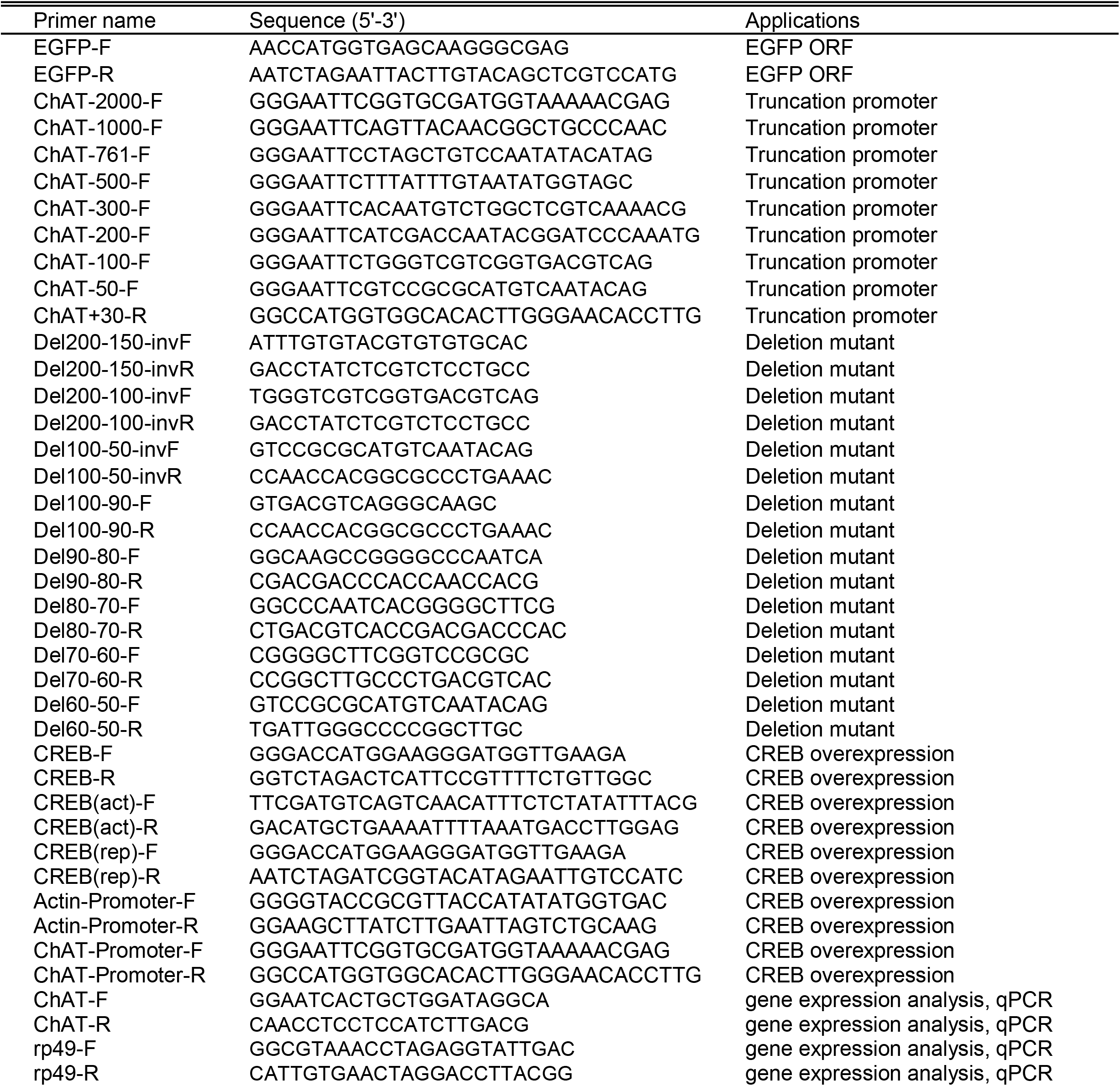
Oligonucleotide sequences of primer used for this study.

**Table 2.**
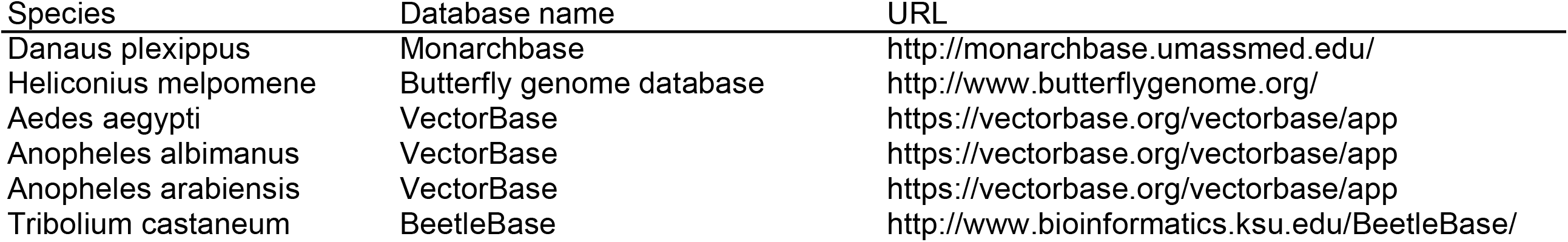
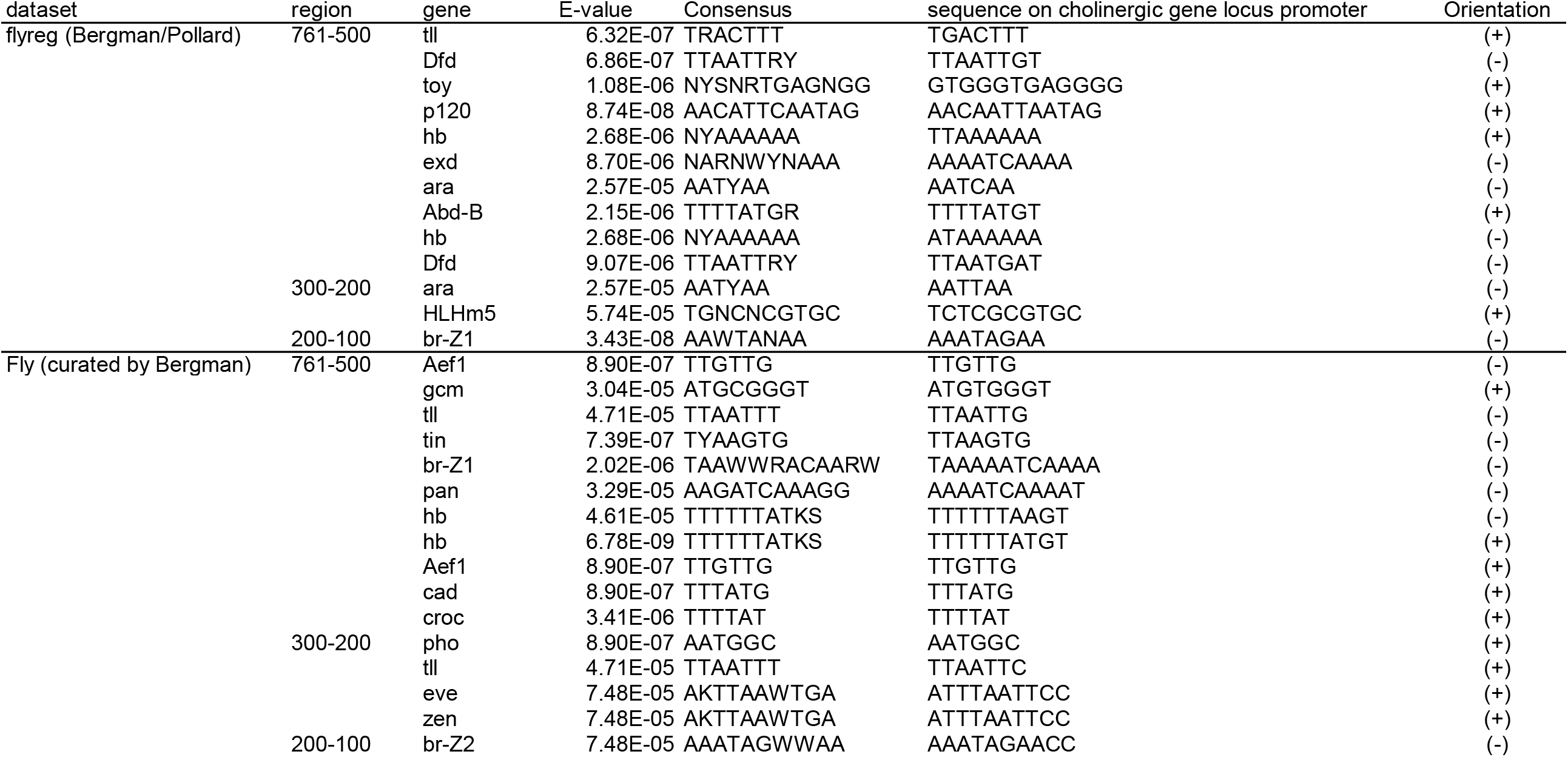
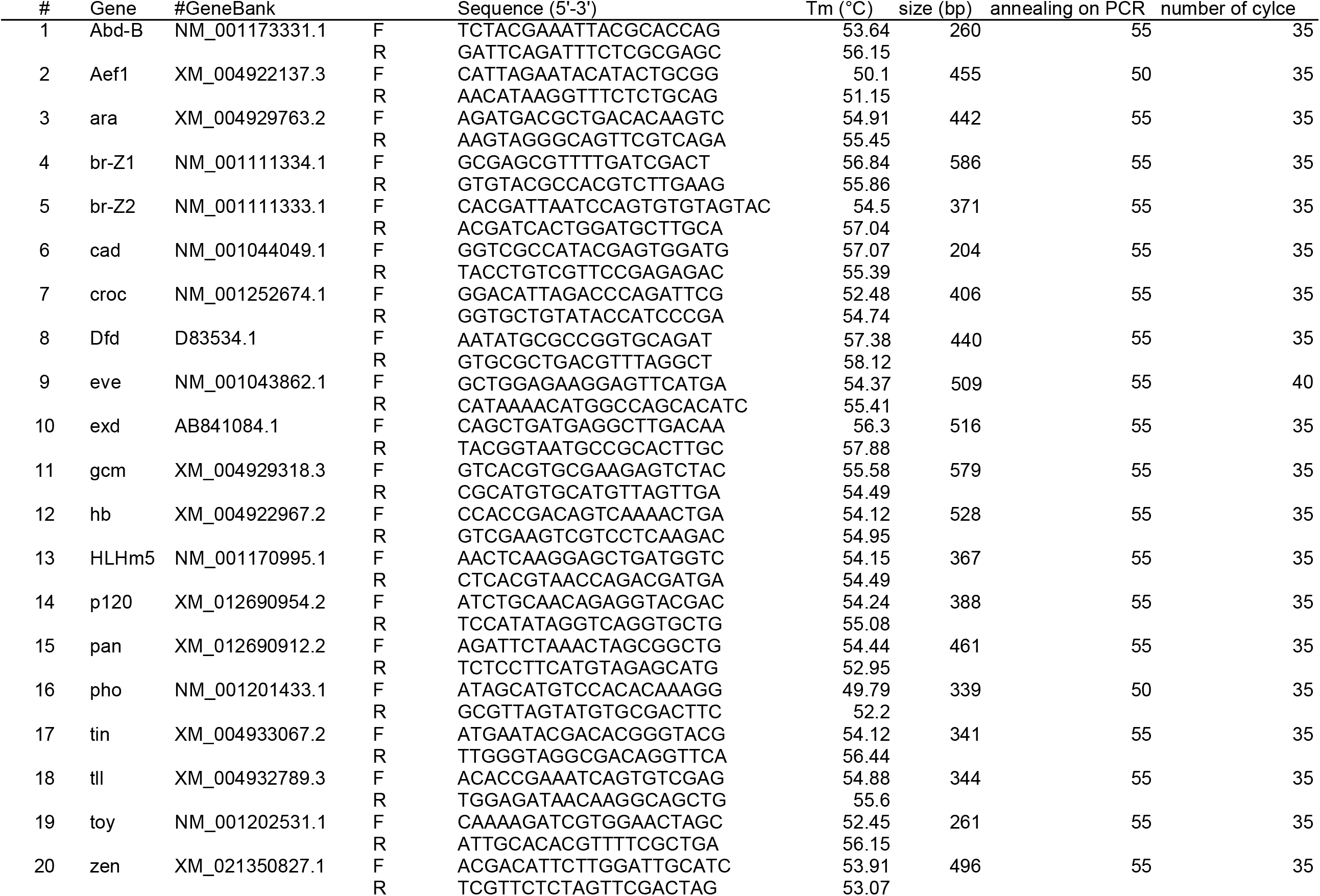
Genomic DNA databases of insect species

## Author contributions

K.B. performed experiments and analyzed data. K.B. and I.S. designed the research and wrote the article.

## Acknowledgements

We thank K. Bui and Y. Ishikawa for comments on thie manucript. This work was supported in part by the grant of Research Institute for Integranted Science from Kanagawa University and the Kanagawa University Grant for Joint Research.

